# Removing independent noise in systems neuroscience data using DeepInterpolation

**DOI:** 10.1101/2020.10.15.341602

**Authors:** Jérôme Lecoq, Michael Oliver, Joshua H. Siegle, Natalia Orlova, Christof Koch

## Abstract

Progress in nearly every scientific discipline is hindered by the presence of independent noise in spatiotemporally structured datasets. Three widespread technologies for measuring neural activity—calcium imaging, extracellular electrophysiology, and fMRI—all operate in domains in which shot noise and/or thermal noise deteriorate the quality of measured physiological signals. Current denoising approaches sacrifice spatial and/or temporal resolution to increase the Signal-to-Noise Ratio of weak neuronal events, leading to missed opportunities for scientific discovery.

Here, we introduce *DeepInterpolation*, a general-purpose denoising algorithm that trains a spatio-temporal nonlinear interpolation model using only noisy samples from the original raw data. Applying DeepInterpolation to *in vivo* two-photon Ca^2+^ imaging yields up to 6 times more segmented neuronal segments with a 15 fold increase in single pixel SNR, uncovering network dynamics at the single-trial level. In extracellular electrophysiology recordings, DeepInterpolation recovered 25% more high-quality spiking units compared to a standard data analysis pipeline. On fMRI datasets, DeepInterpolation increased the SNR of individual voxels 1.6-fold. All these improvements were attained without sacrificing spatial or temporal resolution.

DeepInterpolation could well have a similar impact in other domains for which independent noise is present in experimental data.

## Introduction

Independent noise is a major impediment to any data acquisition effort. Experimental systems neuroscience is acutely impacted given the weakness of neuronal signals. For example, *in vivo* imaging of fluorescent reporters of cellular activity (voltage and calcium) are typically operating in low-photon-count regimes where shot noise largely dominates the recorded signal. Similarly, thermal and shot noise present in electronic circuits independently drown out *in vivo* electrophysiological recordings causing lower sensitivity, impairing the detection of individual spikes. Functional Magnetic Resonance Imaging (fMRI) also suffers from thermal noise present in recording circuits, as relevant changes in Blood-Oxygen-Level Dependent (BOLD) signal are typically just a few percent. The presence of independent noise in all these techniques hampers our ability to measure authentic biological signals, causing irreproducible research or inducing biases related to particular choices of denoising filters^1^ or statistical thresholds^2^.

Often, nearby spatio-temporal samples share signals but are independently affected by noise. This relationship allows the creation of filters to remove noise when applied to the data. Indeed, it is common practice to apply median or Gaussian filters as a first pre-processing step. Sometimes these averaging filters are designed in the Fourier domain to efficiently separate noise from the signal. Although this approach has been successful for many applications, optimal denoising filters can be complicated to build when these data relationships span multiple dimensions (e.g., time and space) or are intrinsically non-linear. Importantly, most of these approaches also impact the signal, causing reductions in spatial or temporal resolution.

The use of machine learning with large datasets permits very complex hierarchical relationships between data points to be learned^3^. In this framework, learned statistical relationships between samples are exploited to reconstruct a noiseless version of the signal, rather than applying filters to remove noise from the signal. This approach has been particularly successful for learning denoising filters or upsampling filters, including for fluorescence imaging^4,5^ but was limited until recently to cases where ground truth is readily available. For example, pre-acquired structural data at various spatial resolutions made it possible to train neural network models to upsample datasets and limit the amount of data necessary to reconstruct a full resolution image^4^. However, when dealing with the types of dynamic signals at the heart of systems neuroscience, denoised datasets or ground truth data are not readily available or may be impossible to obtain.

A recent approach called Noise2Noise^6^ demonstrated that deep neural networks can be trained to perform image restoration without access to any clean ‘ground-truth’ data, with performance comparable or exceeding training using cleaned data. Theoretical work extended this framework to remove pixel-wise independent noise^7,8^. Here, we adapted these recent developments in machine learning to the problem of denoising dynamical signals at the heart of systems neuroscience, thereby building a general-purpose denoising algorithm we call *DeepInterpolation*. Here, we describe and demonstrate the use of *DeepInterpolation* for two-photon *in vivo* Ca^2+^ imaging data, *in vivo* electrophysiological recordings, and whole brain fMRI data. Importantly, our approach is applicable to many experimental fields with minimal modifications.

## Results

The Noise2Noise^6^ study demonstrated that deep neural networks can be trained to perform image restoration without access to any clean ‘ground-truth’ data. At the heart of this approach is the insight that gradients derived from a loss function calculated on noisy samples are still partially aligned toward the correct solution. Provided enough noisy samples are available to train-and the noisy loss is an unbiased estimate of the true loss-the correct denoising filters are still learned^6^. However, unlike the data used in the Noise2Noise publication^6^, systems neuroscience data does not contain pairs of samples with identical signals but different noise.

To address this limitation, we followed a similar approach proposed recently in the Noise2Self^7^ and Noise2Void^8^ frameworks. In the absence of training pairs, we sought to solve an interpolation problem in order to learn the spatio-temporal relationships present in the data. That is, we do not seek to denoise an image but rather to learn the underlying relationship between an entire frame and spatio-temporally nearby samples by optimizing the reconstruction loss calculated on noisy instances of the center frame. Second, the noise present in the target (center) samples needs to be independent from the input (adjacent) samples; otherwise our relationship model would overfit the noise we seek to remove. We eliminated any chance of overfitting by (1) omitting the target center frame from the input, and (2) presenting training samples only once during training. While this noisy center frame is the most informative about its own value, it serves the key function of the target during training. During inference, the dataset is simply streamed through the trained network to reconstruct a noiseless version of the signal. The quality of the reconstruction depends on whether enough information about the center frame was present in spatio-temporally nearby samples and if it was appropriately learned by the training procedure. Given the extensive leverage of complex relationships present in the data, spanning multiple spatio-temporal dimensions, we will refer to this general framework as ***DeepInterpolation***.

### Application of DeepInterpolation to *in vivo* two-photon Ca^2+^ imaging

For two-photon imaging, we constructed our denoising network following simple principles: First, given the presence of neuropil signals distributed throughout the background, a single pixel can share information with other pixels throughout the entire field of view. Second, the decay dynamics of calcium-dependent fluorescent indicators (GCaMP6f τ_peak_ = 80 +/− 35 ms; τ_1/2_ = 400 +/− 41 ms^9^) suggest frames as far as 1 second away can carry meaningful information. Therefore, we chose to train a neuronal network to reconstruct a center frame from the data present in N_pre_ prior frames and N_post_ subsequent frames. The center frame was omitted from the input so as to not provide any information about the independent pixel-wise noise due to shot noise (**Fig. 1A**). The final value of the meta-parameter N_pre_ and N_post_ were selected by comparing the validation loss during training (**Fig. 1B**, see **Methods** and **Appendix I** for details on loss calculation). Our denoising network utilized a UNet inspired encoder-decoder architecture with 2D convolutional layers^10,11^.

**Fig. 1.**
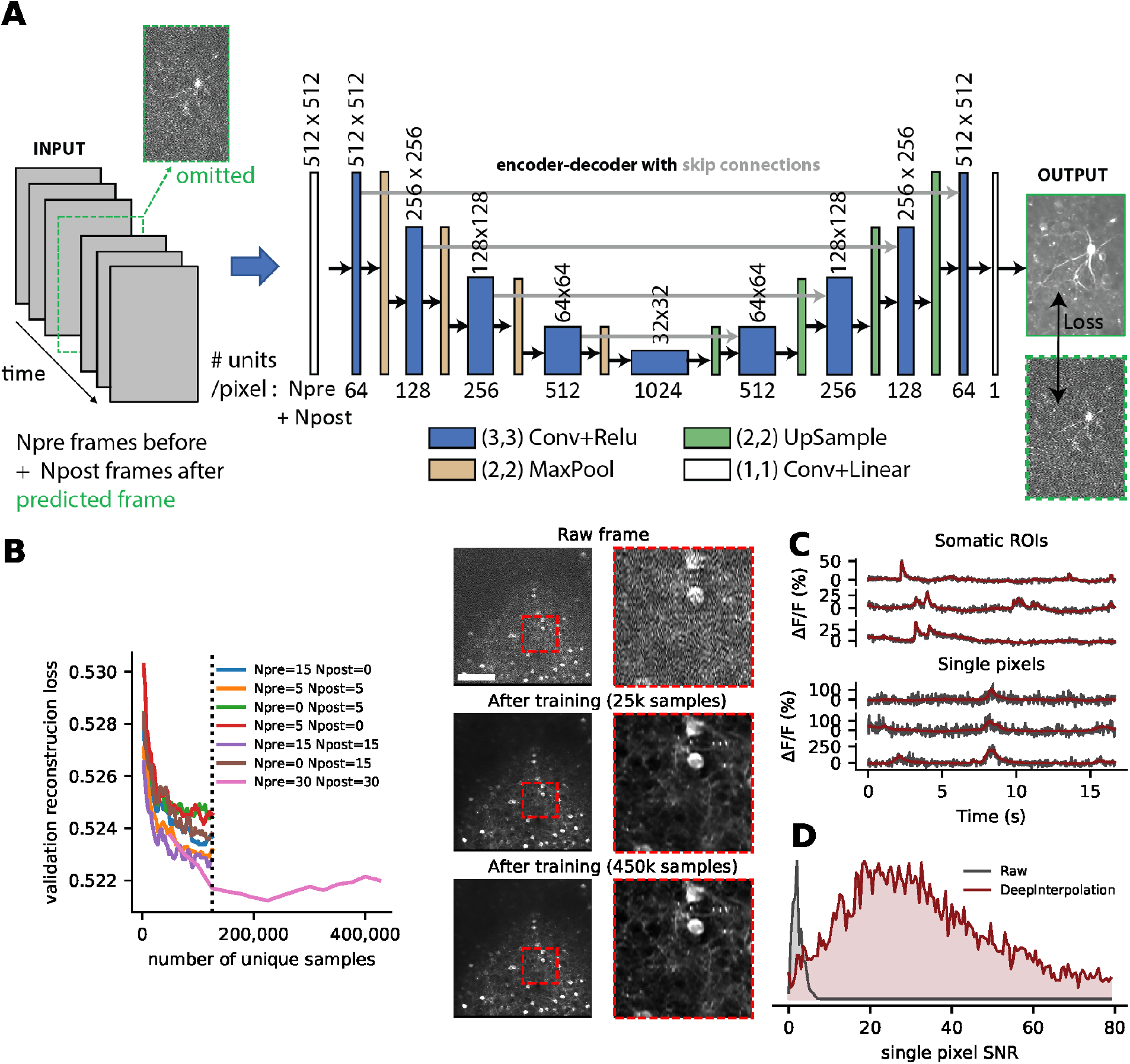
Training DeepInterpolation networks for denoising two-photon Ca^2+^ imaging. **(A)** An encoder-decoder deep network with skip connections was trained to denoise two-photon imaging data. The training procedure minimized the L1 loss between the predicted frame and a raw frame contaminated with shot noise. The encoder-decoder network utilized N_pre_ and N_post_ frames, acquired at 30 Hz, respectively, before and after the target frame (labelled in green) to predict the central, target frame. (which was omitted from the input). **(B)** To validate training performance, the validation loss was monitored for convergence as well as the quality of the image reconstruction throughout training (see inset). Y axis is the mean absolute difference between a predicted frame and a noisy sample across 2500 samples. Individual data samples were z-scored using a single estimate of mean and standard deviation per movie (see Methods). Different combinations of (N_pre_, N_post_) values were tested during training. Dashed vertical line indicates early stopping (due to the extreme computational demands during training) to evaluate the best set of parameters. We continued training the model with N_pre_=30 frames and N_post_=30 frames and used this set of parameters for the rest of this study. Right panels illustrate the denoising performance of this model compared to a single raw frame (top). Notice the improved contrast between 25,000 and 450,000 samples training: the smallest neuronal compartments are more clearly visible in the latter. Scale bar is 100 μm. Each red inset is 100 μm. (**C**) Six example traces extracted from a held-out denoised movie before (black) and after (red) denoising with DeepInterpolation. The top three traces are extracted from a somatic ROI, while the bottom three traces are extracted from a single pixel. (**D**) Distribution of SNR (mean over standard deviation, see Methods) for 10,000 voxels (randomly selected across N=19 denoised held-out test movies) before and after DeepInterpolation.

The Allen Brain Observatory datasets (http://observatory.brain-map.org/visualcoding) offered a unique opportunity to train these denoising networks as we had collected more than 1,300 hours of highly standardized and curated datasets^12^. For each Gcamp6 reporter line (Ai93, Ai148), we had access to more than 100 million frames carefully curated with standard Quality Control (QC) metrics. We leveraged this database by presenting each training sample only once during the entire training process, completely eliminating any chance of noise overfitting.

Within the limits of our computational infrastructure, we showed that using a larger number of both prior and subsequent frames as input yielded smaller final validation reconstruction losses (see **Fig. 1B**). Training required 225,000 samples pulled randomly from 1144 separate one hour long experiments to stop improving (see **Fig. 1B**). Since the loss is dominated by the independent noise present in the target image, the reconstruction loss could not converge to zero. This was supported by simulation of ideal reconstruction losses with known ground truth (**Supp. Fig. 1**). In fact, even small improvement in the loss related to visible improvements in the reconstruction quality of the signal (**Fig. 1B**).

Applying this trained network to denoise held-out datasets (**Supp. Video 1, 2**) yielded remarkable improvements in imaging quality. The same trained network generalized well to many movie examples (see 4 examples on **Supp. Video 2**). Shot noise was visibly eliminated from the reconstruction (see single frames in **Fig. 1C** and associated calcium trace) while calcium dynamics were preserved, yielding a 15 fold increase in SNR (mean raw pixel SNR = 2.4 +/− 0.01, mean pixel SNR after DeepInterpolation = 37.2 +/− 0.2, N=9966 pixels). While the movies we used during training were motion corrected, a small amount of motion remained and close inspection of the reconstructions showed that the algorithm also corrected these remaining motion artifacts (**Supp. Video 1, 2**).

We compared our denoising approach with a recent algorithm called Penalized Matrix Decomposition (PMD)^13^. While PMD properly reconstructed somatic activities, it rejected most of the variance present in background pixels, contrary to DeepInterpolation (see **Supp. Video 3**). A key underlying assumption of PMD is that motion artifacts are absent from the data. Contrary to DeepInterpolation movies, we observed clear artifacts in movies denoised with PMD due to small residual motion artifacts (see arrow in **Supp. Video 3**).

To compare these two approaches given this limitation, we created entirely simulated datasets devoid of motion artifacts, for which we had knowledge of the underlying ground truth. For this, we used a recent approach called *in silico* Neural Anatomy and Optical Microscopy (NAOMi)^14^. NAOMi combines a detailed anatomical model of a volume of tissue with models of light propagation and laser scanning so as to generate realistic Ca^2+^ imaging datasets (see **Methods**). Both DeepInterpolation and PMD performed well with datasets generated by NAOMi (see **Supp. Fig. 2 B,C** and **Supp. Video 4**). Because DeepInterpolation used information from nearby frames, it provided visibly smoother calcium traces with reconstruction losses 1.2 to 2 times smaller than with PMD (see **Supp. Fig. 2**).

Having established that DeepInterpolation compares favorably to the state of the art, we investigated how removing shot noise impacts our ability to analyze the activity of neuronal compartments.

### Segmentation of active compartments during Ca^2+^ imaging is transformed by DeepInterpolation

We first sought to evaluate the impact of DeepInterpolation on the segmentation process itself. Existing approaches to calcium imaging segmentation rely on the analysis of correlation between pixels to create ROIs^15,16^. After denoising, single pixel pairwise correlations greatly increased from near zero (averaged Pearson correlation = 9.0*10^−4^ +/− 2.3*10^−6^ s.e.m, *n* = 4*10^8^ pairs of pixels across 4 experiments) to a significant positive value (averaged Pearson correlation = 0.10 +/− 1.210^−5^ s.e.m, *n* = 4e8 pairs of pixels across 4 experiments, KS test comparing raw with DeepInterpolation: *p* = 9*10^−71^, *n* = 1,000 pixels randomly sub-selected, **Supp. Fig 3**). We expected that this increase in averaged correlation would improve the quality and the number of segmented regions. To that end, we leveraged the sparse mode algorithm available in Suite2p^16^.

The improvement in correlation structure benefited this algorithm considerably. Suite2p extracted a large number of additional active compartments in the field of view, including small sets of pixels covered by axons or dendrites perpendicular to the imaging plane (**Fig. 2A**). In the absence of shot noise and single frame motion artifacts, cell filters were surprisingly well defined (**Fig. 2B**). Automatically extracted ROIs showed precise details of each compartment, sometimes encompassing dendrites or axons connected to a local soma. In some cases, active attached buttons were clearly visible in the extracted filter of an horizontal dendrite (**Fig. 2B**). In many movies, a large number of smaller, punctated compartments - likely a mixture of horizontal sections of apical dendrites or axons - were detected, each with clear calcium transients (**Fig. 2C**).

**Fig. 2.**
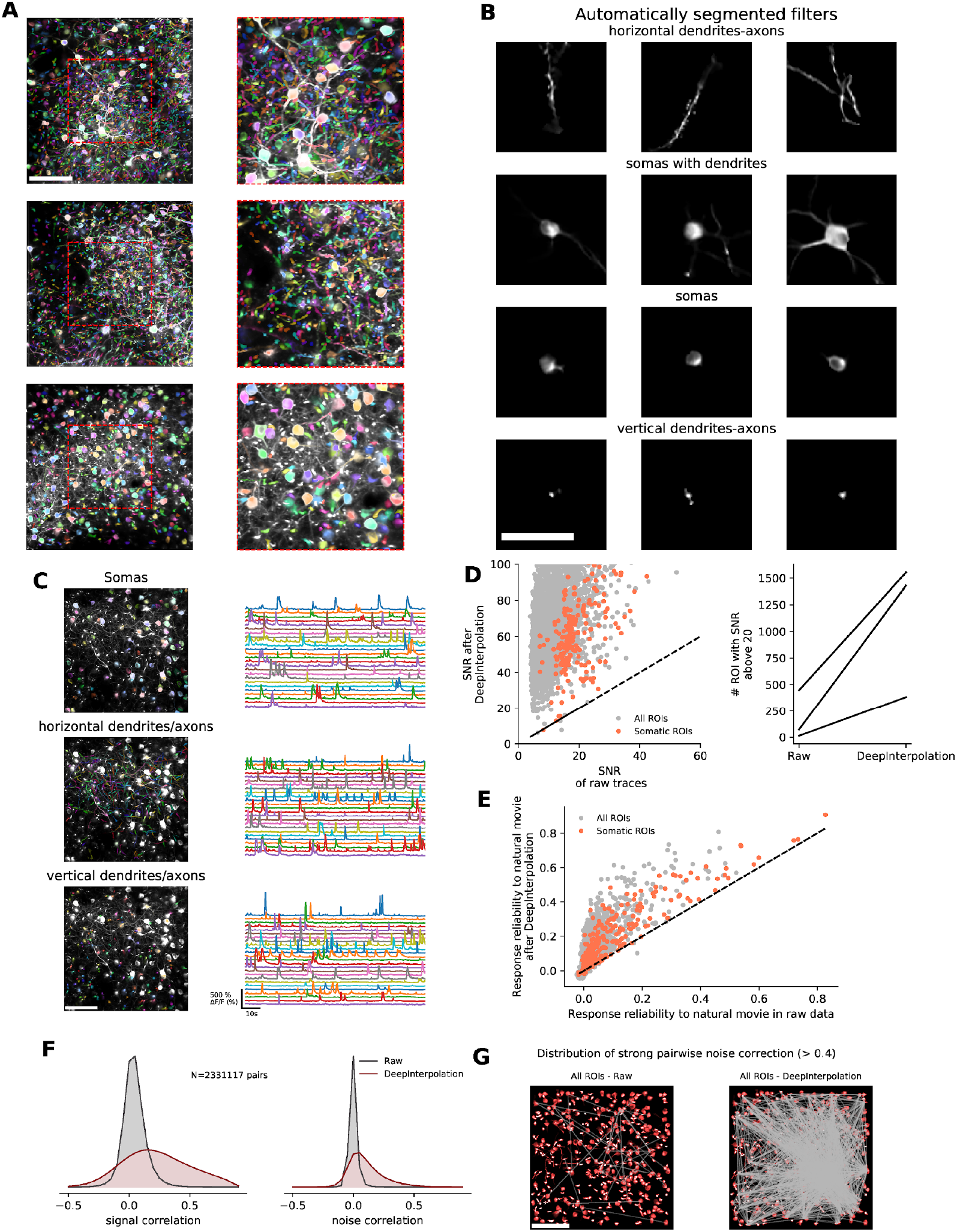
Applying DeepInterpolation to Ca^2+^ imaging reveals many more ROIs and rich network physiology across small and large neuronal compartments. **(A)** Three examples showing overlaid colored segmentation masks on top of a maximum projection image of calcium imaging. Improved correlation structure (see **Supp. Fig 4**) allows the detection of a large number of ROIs using the “sparsity” segmentation mode in Suite2p^16^. Red dashed boxes show zoomed-in view over smaller segmented compartments. Apical and proximal dendrites are cleanly segmented, regardless of whether they were imaged along their length or through a short vertical section. Scale bar is 100 μm. **(B)** Example individual weighted segmented filters showcasing dendrites, isolated somas, somas with attached dendrites, and axons and small sections of dendrites/axons. Scale bar is 50 μm **(C)** Manually sorted ROIs from one experiment showcasing the calcium traces extracted from each individual type of neuronal compartments (from top to bottom - cell body, horizontal processes, processes perpendicular to imaging plane). After DeepInterpolation, calcium events are recorded with high SNR in all three compartment types, regardless of their size. (**D, left**) Quantification of SNR for all detected ROIs (gray dots) and somatic ROIs (red dots) with and without DeepInterpolation. The majority of ROIs have an SNR above 20 after DeepInterpolation. Dashed line represents the identity line. (**D, right**) Comparison of the number of ROIs with an SNR above 20 with and without DeepInterpolation. DeepInterpolation allows to reliably record the activity of more than a thousand compartments in a small (400 μm)^2^ field of view (*n* = 3 experiments). (**E**) Quantification of the response reliability of individual ROIs across 10 trials of a natural movie visual stimulus. Both somatic and non-somatic ROIs see increased response reliability to a visual stimulus. Dashed line represents the identity line. **(F, left)** signal correlation (average correlation coefficient between the average temporal response of a pair of neurons) for all pairs of ROI in (**D**) for both raw and denoised traces. **(F, right)** noise correlation (average correlation coefficient at all time points of the mean-subtracted temporal response of a pair of neurons) for all pairs of ROIs in (**D**). (**G**) For an example experiment, pairs of ROIs with high noise correlation (>0.4) are connected with a straight line for both original two-photon data and after DeepInterpolation. Scale bar is 100 μm.

Although those numerous compartments were not detected in the original movie, we applied the weighted masks to both denoised and raw movies to quantify the improvement in SNR of the associated temporal trace. ROIs associated with somatic compartments typically yielded a mean SNR of 21.0 +/− 0.5 (s.e.m, *n* = 240 in 3 experiment) in the original movie (**Fig. 2D**); after denoising, the majority of those ROIs had an SNR above 40 (mean 73.8 +/− 0.5). Similarly, the mean SNR across all compartments, including the smallest apical dendrites, went from a mean of 13.2 +/− 0.1 (*n* = 3385 in 3 experiments) to 73.8 +/− 0.5. Assuming that a single spike causes a calcium rise of about 10%^9^, an SNR above 20 is necessary for those events to be twice as large as the noise. Although this is a necessary but not sufficient condition, this suggests that removing shot noise will increase our capability to detect the presence of single spikes.

The average number of active detected ROIs with SNR above 20 went from 178 +/− 135 (s.e.m., in 3 experiments) to 1122 +/− 371 in a 400×400 um^2^ field of view (**Fig. 2D**). That is, as much as six times more neuronal compartments are now available from a single movie for analysis following DeepInterpolation, thanks to a more reliable segmentation process.

### DeepInterpolation improves the analysis of correlated Ca^2+^ imaging activity

To illustrate this new opportunity, we analyzed the response of all compartments to 10 repeats of a natural movie stimulation. We found clear examples of large and smaller responses to natural movies in both somatic and non-somatic compartments (**Supp. Fig. 4A**). The trial by trial response reliability (see **Methods**) increased substantially from raw traces to traces after DeepInterpolation (see, **Fig. 2E**), both for somatics ROIs (raw: mean 0.10 +/− 0.01, DeepInterpolation: mean 0.20 +/− 0.01; s.e.m, *n* = 240 ROIs in *n* = 3 experiments) and across all ROIs (raw: mean 0.029 +/− 0.001, DeepInterpolation: mean 0.113 +/− 0.002; s.e.m, *n* = 3385 ROIs in *n* = 3 experiments). When considering the distribution of these responses across a single field of view, both somatic and non-somatic compartments had a similar distribution for their preferred movie frame, the maximum DF/F response at this prefered stimulus, as well as the reliability of their response to natural movies (see **Supp. Fig. 4B, C, D**).

Neuronal representations are inherently noisy even when sensory stimuli and the behavioral state are kept constant. This key property of neuronal networks has fueled a large number of studies on the relationship between the averaged signal representation in individual neurons and the trial by trial fluctuation of these responses^17,18^. Experimental noise directly impacts the measure of neuronal relationships as it modulates trial-by-trial responses.

For each pair of ROIs in a single experiment, we extracted their shared signal and noise correlation (see **Methods**). As expected, we found that both signal as well as noise correlation increased after denoising (signal: from 0.06725 ± 0.00006 to 0.25 ± 0.0001; noise from 0.02240 ± 0.00002 to 0.12351 ± 0.00008; both for 2,329,554 pairs of ROIs from three experiments; **Fig. 2F**). This result was not just due to the improved segmentation as it was preserved using ROIs only detected on raw noisy data (see **Supp. Fig. 5**).

To further illustrate how these improved measures of neuronal fluctuations impacted the analysis of functional interactions, we compared the spatial distribution of pairwise correlations between neurons during natural movie stimulation. DeepInterpolation largely increased the number of strong pairwise interactions for both somatic and non somatic pairs (see **Fig. 2G**, all ROIs: 43 to 1721 pairs). These correlated pairs of units were almost invisible in the original data, illustrating the improved biological insight gained through denoising.

### Application of DeepInterpolation to *in vivo* electrophysiological recordings

Electrophysiological recordings from high-density silicon probes have similar characteristics to two-photon imaging movies: information is shared across nearby pixels (electrodes), as well as across time. We therefore reasoned that a similar architecture should perform well for denoising electrophysiology data, in particular for removing strictly independent thermal and shot noise present in the recordings. For training data, we used Neuropixels data that were similarly standardized and curated as the two-photon recordings^19^.

Because a single action potential lasts ~1 ms, we constructed our interpolating filter to predict the voltage profile at any given moment from 30 preceding and 30 following time steps acquired at 30 kHz. Contrary to two-photon imaging, we found that omitting one sample before and after the center sample better rejected the background noise while keeping the signal reconstruction of the highest quality. The architecture of the network was similar to the one used for two-photon imaging, except that the input layer was reshaped to match the approximate layout of the Neuropixels recording sites (**Fig. 3A**). Because electrophysiological recordings have a 1,000-fold higher sampling rate than two-photon imaging, we had access to essentially 3 orders of magnitude more training data per experiment and could train the network using only 3 experiments in total.

After inference, a qualitative inspection of the recordings showcased excellent noise rejection (**Fig. 3B**). We compared spike profiles before and after denoising and found that the spike waveforms were more clearly visible in the denoised data, especially for units with lower amplitude spikes (**Fig. 3B**, inset; **Fig. 3C**). The residual of our reconstruction showed weak spike structure but spectral analysis revealed a mostly flat frequency distribution (**Supp. Fig. 6A**). Importantly, the overall shape (amplitude and decay time) of individual action potentials was preserved, while the median channel RMS noise was decreased 1.7-fold (6.96 μV after denoising, see **Fig. 3D**). We ran a state-of-the-art spike sorting algorithm (Kilosort2, version downloaded April 8, 2019) and removed any detected units with waveforms not likely to be associated with action potentials. After denoising, we found 25.5 ± 14.5% more high-quality units per probe (ISI violations score < 0.5, amplitude cutoff < 0.1, presence ratio > 0.95; see Methods for details). The number of detected units was higher after DeepInterpolation regardless of the quality metric thresholds that were chosen (**Supp. Fig. 6B**). The majority of additional units had low-amplitude waveforms, with the number of units with waveform amplitudes above 75 μV remaining roughly the same (1532 before vs. 1511 after), while the number with amplitudes below 75 μV increased dramatically (381 before vs. 910 after) (**Fig. 3E**). For units that were clearly matched before and after denoising (at least 70% overlapping spikes), average SNR (defined as the ratio of the waveform amplitude to the RMS noise on the peak channel) increased from 11.2 to 14.5 (**Fig. 3F**).

**Fig. 3.**
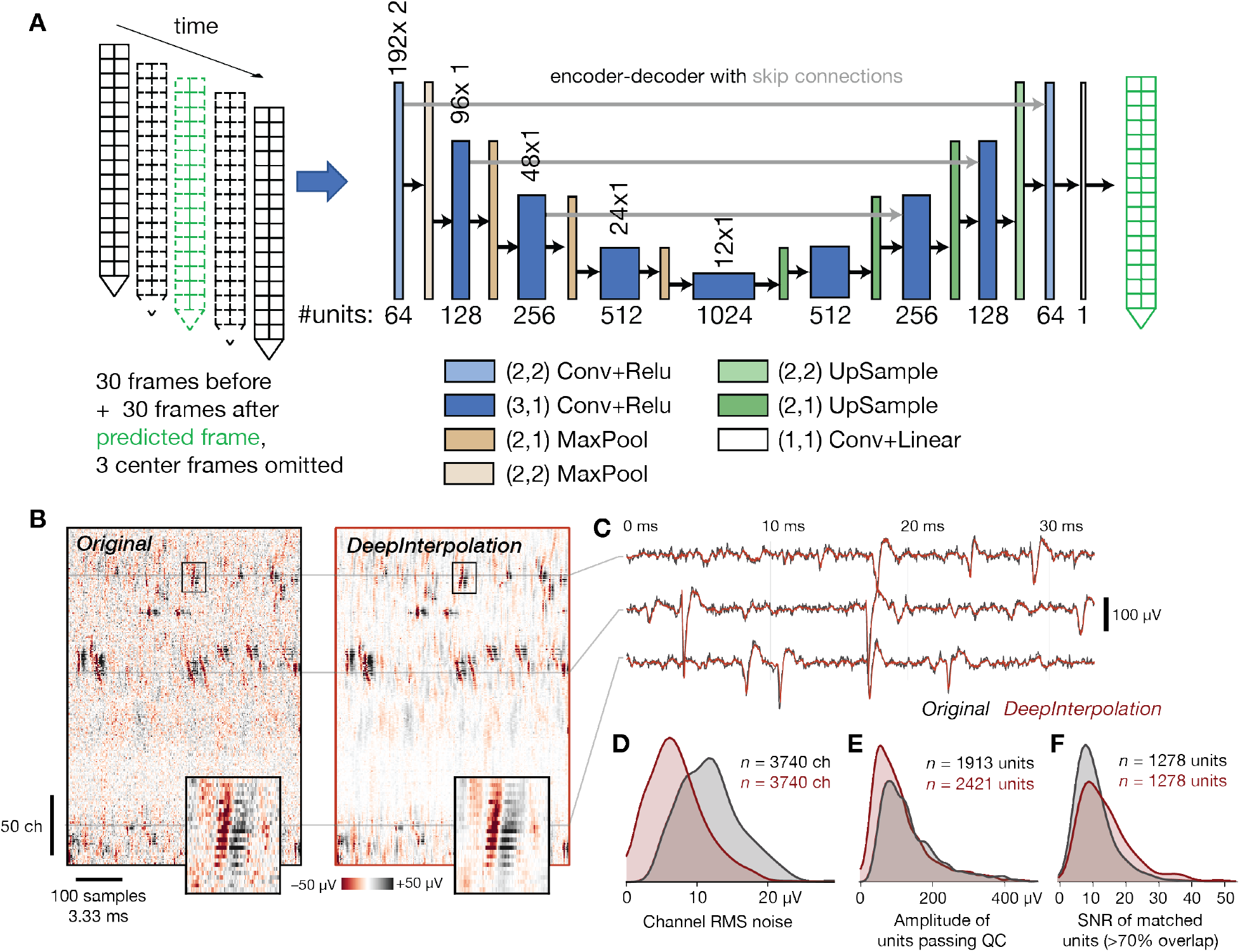
Applying DeepInterpolation to electrophysiological recordings decreases background noise and yields more neuronal units. (**A**) Structure of the DeepInterpolation encoder-decoder network for electrophysiological data recorded from a Neuropixels silicon probe, with two rows of 192 electrodes each, spaced 20 μm apart. Data is acquired at 30 kHz. (**B**) Side-by-side comparison of original (left) and denoised (right) data, plotted as a 2D heatmap with time on the horizontal axis and channels along the vertical axis. Insets show a close-up of a single action potential across 22 contiguous channels (the box is 3.6 ms wide). (**C**) Denoised time series (red) overlaid on the original time series (gray) for three channels from panel B. Histogram of RMS noise for all channels from 10 experiments, before and after applying DeepInterpolation. **(E)** Histogram of waveform amplitudes for all high-quality units from 10 experiments. **(F)** Histogram of waveform SNRs for all units that were matched before and after denoising.

By searching for spatiotemporally overlapping spikes, we determined that 6.2 ± 2.9% of units in the original recording were no longer detected after applying DeepInterpolation (<50% matching spikes). However, this was counterbalanced by the addition of 20.2 ± 4.8% units that had fewer than 50% matching spikes in the original data (example unit in **Supp. Fig. 6C**). This is less than the 25.5% increase cited above, because it accounts for a minority of units that were detected but merged together in the original data. To validate that these novel units were likely to correspond to actual neuronal compartments, we analyzed their responses to natural movie stimulation, using the same movie as in the two-photon imaging study. We found approximately 9% more reliably visually modulated units in the visual cortex after denoising (**Supp. Fig. 6D**), and found clear examples of stimulus-modulated units that were previously undetected (**Supp. Fig. 6E**).

Applying DeepInterpolation to extracellular electrophysiology data increased the unit yield across all brain regions tested, including visual cortex, hippocampus, and thalamus (**Supp. Fig. 7A**). Additional units tended to have firing rates in the middle of each region's average range and waveform amplitudes below 100 μV (**Supp. Fig. 7B**). In all regions except for dentate gyrus, DeepInterpolation preserved the original waveform amplitude, while detecting more low-amplitude units. The downward shift in waveform amplitude observed in dentate gyrus suggests that it may be necessary to provide additional training data from this region in order to optimize performance of the denoising step. In the visual cortex, additional units were primarily “regular-spiking” (waveform peak-to-trough duration > 0.4 ms), rather than “fast-spiking.” This suggests that these may be previously undetectable regular-spiking interneurons with small cell bodies; however, applying DeepInterpolation to datasets with ground truth cell type labels obtained via optotagging^20^ will be required to confirm this hypothesis.

### Application of DeepInterpolation to functional Magnetic Resonance Imaging *(fMRI)*

We sought to evaluate how DeepInterpolation could help the analysis of volumetric datasets like fMRI. fMRI is very noisy as the BOLD response is typically just a 1-2 % change of the total signal amplitude^21^. Thermal noise is present in the electrical circuit used for receiving the MR signal. A typical fMRI processing pipeline involves averaging nearby pixels and successive trials to increase the SNR^22,23^. We reasoned that our approach could replace smoothing kernels with more optimal local interpolation functions without sacrificing spatial or temporal resolution.

Because of the low sampling rate (0.3 to 0.5 Hz), a single fMRI recording session could only provide several hundreds full brain volumes, while each volume typically contains up to 10 millions voxels. To augment our training datasets, we decided to alter our approach: Instead of learning a full brain interpolation model, we sought to train a more local interpolation function.

To reconstruct the value of a brain sub-volume, we fed a neural network with a consecutive temporal series of a local volume of 7 × 7 × 7 voxels. The entire target volume was omitted from the input (**Fig. 4A**). To allow our interpolation network to be robust to edge conditions, input volumes on the edge of the volume were fed with zeros for missing values. For inference, we convolved the denoising network through all voxels of the volume, across both space and time, using only the center pixel of the output reconstructed volume to avoid any potential volume boundaries artifacts.

**Fig. 4.**
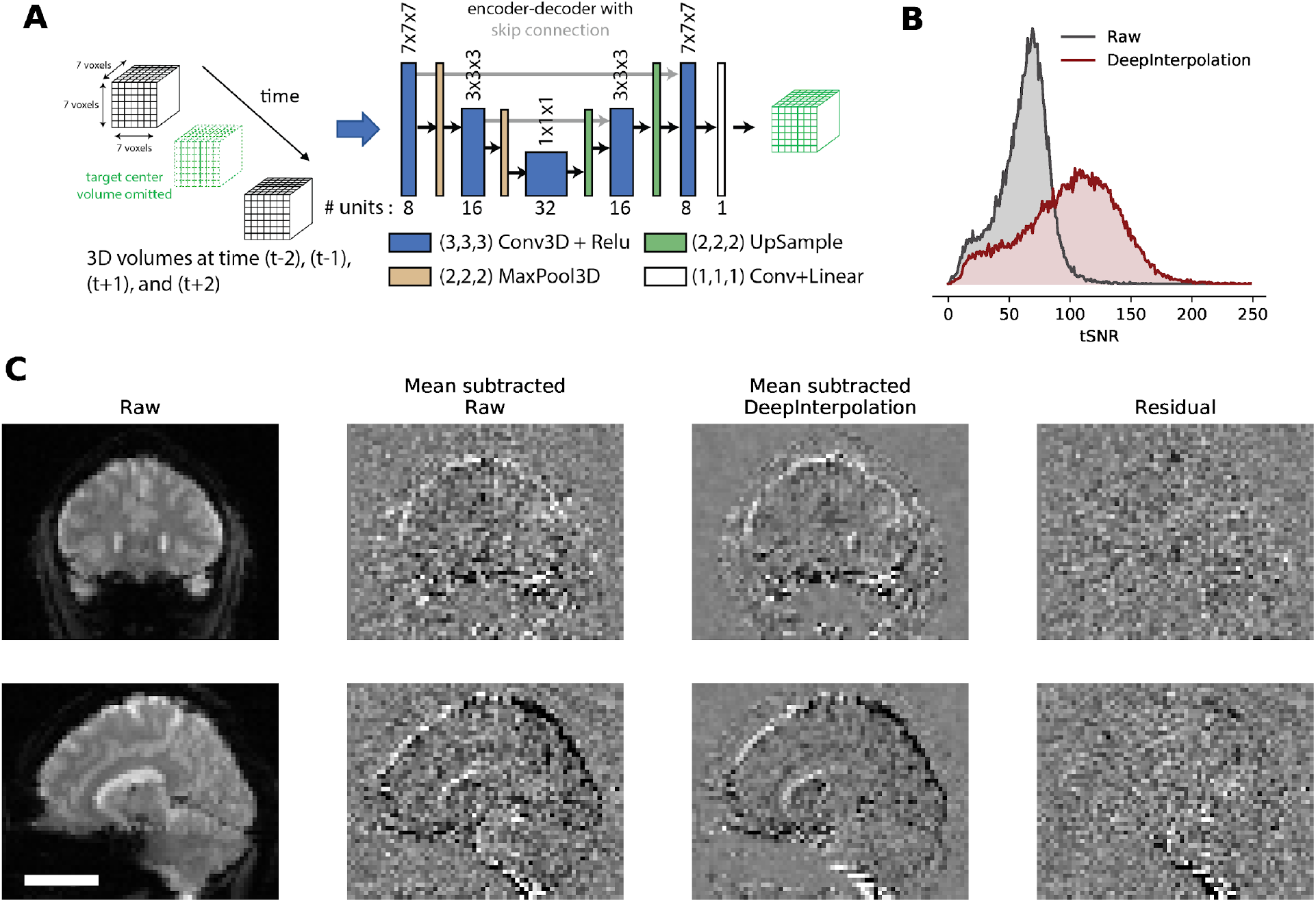
Applying DeepInterpolation to fMRI removes thermal noise. (**A**) Structure of the DeepInterpolation encoder-decoder network for fMRI data. Instead of predicting a whole brain volume at once, the network reconstructs a local 7×7×7 cube of voxels. (**B**) tSNR (see Methods) for 10,000 voxels randomly distributed in the brain volume in raw data and after DeepInterpolation. (**C**) Exemplar reconstruction of a single fMRI volume. First row is a coronal section while the second row is a sagittal section through a human brain. Scale bar is 5 cm. In the second column, the temporal mean of 3D scan was removed to better illustrate the presence of thermal noise in the raw data. The local denoising network was processed throughout the whole 3D scan for denoising. The impact of DeepInterpolation on thermal noise is illustrated in the 3rd column. The 4th column shows the residual of the denoising process. Notice how the residual captured independent noise without any signal structure. Occasional large blood vessels are visible in the residual (see bottom).

We trained a single denoising network using 140 fMRI datasets acquired across 5 subjects or 1.2 billion samples. Each training sample was presented only once to avoid any chance of overfitting. After training, we applied the network to held-out test data, showing excellent denoising performance (**Fig. 4C**).

We extracted the temporal traces from individual voxels and found that tSNR (see **Methods**) increased from 61.6 ± 20.6 to 100.40 ± 38.7 (*n* = 100,001 voxels). We noticed that the background noise around the head was clearly excluded and only present in the residual image (**Fig. 4C** **and Supp. Video 5**). Surrounding soft tissues became clearly visible after denoising (see **Fig. 4C,** Raw data: background voxel std = 7.59 ± 0.01 s.e.m; brain voxels std = 15.95 ± 0.08 s.em., *n* = 10,000 voxels; DeepInterpolation data: background std = 2.24 ± 0.01 s.em., *n* = 10,000; brain voxels std = 9.72 ± 0.05 s.em., *n* = 10,000 voxels; Residual: background std = 7.91 ± 0.01; brain voxels std = 15.79 ± 0.08, *n* = 10,000 voxels). The residual movie showed no visible structure except for occasional blood vessels, corrected motion artifacts and head mounting hardware (**Fig. 4C, Supp. Video 7**). We compared movies extracted with DeepInterpolation to fully pre-processed movies that included a Gaussian denoising kernel (**Supp. Video 6**): the original 3D resolution was fully maintained in DeepInterpolation while the processed data with Gaussian kernel clearly diffused the voxel dynamic across all nearby voxels.

## Discussion

Here, we demonstrated a methodology, *DeepInterpolation*, related to the approach outlined in the *Noise2Self^7^* and *Noise2Void*^8^ framework to reconstruct noiseless versions of dynamic datasets, without any requirement for ground truth. While we share the key separation of noise and signal with these techniques, we developed our framework independently. As a result, several key differences are notable. First, instead of working with single fluorescence frames with pixel-wise omissions, we adapted our approach to complex multi-dimensional biological datasets at the heart of systems neuroscience. Second, we trained our models on large databases, demonstrating the impact of richer denoising models on existing neuroscience scientific data and data workflows like cell segmentations or unit detection. Third, we contributed practical solutions: network topology, training regimen and a validation process to extract complex multi-dimensional models.

When applied to two-photon Ca^2+^ imaging datasets, DeepInterpolation increased the SNR of single pixels 15-fold. On electrophysiological data, with little architectural change, DeepInterpolation uncovered 25% more high quality neuronal units. Similarly, applying DeepInterpolation led to a 1.6-fold increase in the SNR of fMRI voxels, effectively removing thermal noise from the recordings. The successful application of DeepInterpolation to these three experimental data modalities supports the general relevance of our method.

Given the increased leverage of the information present between samples, we anticipate DeepInterpolation to be instrumental for the development of large scale voltage-imaging of neuronal activity^24,25^ as well as facilitate high-speed fMRI where thermal noise dominates the BOLD signal. Importantly, the pixel-wise framework we used for fMRI increased our training dataset, offering the opportunity to apply the same methodology to many different modalities, regardless of whether or not vast quantities of training data are available. Our approach should permit scientists in a variety of fields to re-analyze their existing datasets but now with independent noise removed.

Speeding up and/or parallelizing the inference step could facilitate real-time denoising of spatiotemporal signals. This could enable a process whereby a short dataset would be acquired at the onset of any one experiment, similar to a calibration routine. This dataset could then be used to fine-tune a pre-trained denoiser that removes independent noise. Whether such training needs to be specific for each instrument, each type of instrument, each brain area, each transgenic mouse line or even each subject remains a topic for future research.

We here demonstrated a general purpose denoising methodology to reconstruct dynamic signals at the heart of systems neuroscience without the contamination of independent noise. Applying DeepInterpolation to Ca^2+^ imaging, electrophysiological recordings and fMRI showed its wide impacts on SNR and its ability to uncover previous hidden neuronal dynamics. We anticipate that future work in all 3 modalities we presented here will revise key subsequent processing steps to better leverage clearer correlation structures now measured in the data.

## Acknowledgement

We thank the Allen Institute founder, Paul G. Allen, for his vision, encouragement, and support. We thank Alex Song from the Tank Laboratory for help with NAOMi and for providing early access to the codebase that he created with Adam Charles. We thank Liam Paninski, Saveliy Yusufov, and Ian August Kinsella for sharing code and support for applying PMD onto comparative datasets. We thank Daniel Kapner for help with two-photon segmentation as well as for supportive discussions. We thank James Philip Thomson from the Schnitzer laboratory for discussions related to denoising two-photon movies. We thank Anton Arkhipov and Nathan Gouwens for fruitful discussions and for supporting early use of DeepInterpolation toward analyzing two-photon datasets. We thank Mark Schnitzer and Biafra Ahanonu for continuous support and evaluating early application of DeepInterpolation to one-photon datasets. We thank Todd Peterson, Myra Imanaka, Steve Lawrenz, Sarah Naylor, Gabe Ocker, Michael Buice, Jack Waters, Peter Ledochowitsch and Hongkui Zeng for encouragement and support. We thank Doug Ollerenshaw and Nathan Gouwens for sharing code to standardize figure generation.

## Author Contributions

J.L. initiated, conceptualized and directed the project. J.L. and M.O. conceptualized the DeepInterpolation framework. J.L. wrote all training and inference code with guidance from M.O. J.L. performed all two-photon and fMRI analysis and ran DeepInterpolation on electrophysiological data. J.H.S and J.L. iterated the electrophysiological denoising process. J.H.S. performed all analysis on electrophysiological data. C.K. guided the characterization of all denoising performance. N.O. ran the two-photon ground truth simulation. J.L., M.O., J.H.S and C.K wrote the paper.

**Correspondence and requests** for materials should be addressed to J.L.

## Supplementary Video Table

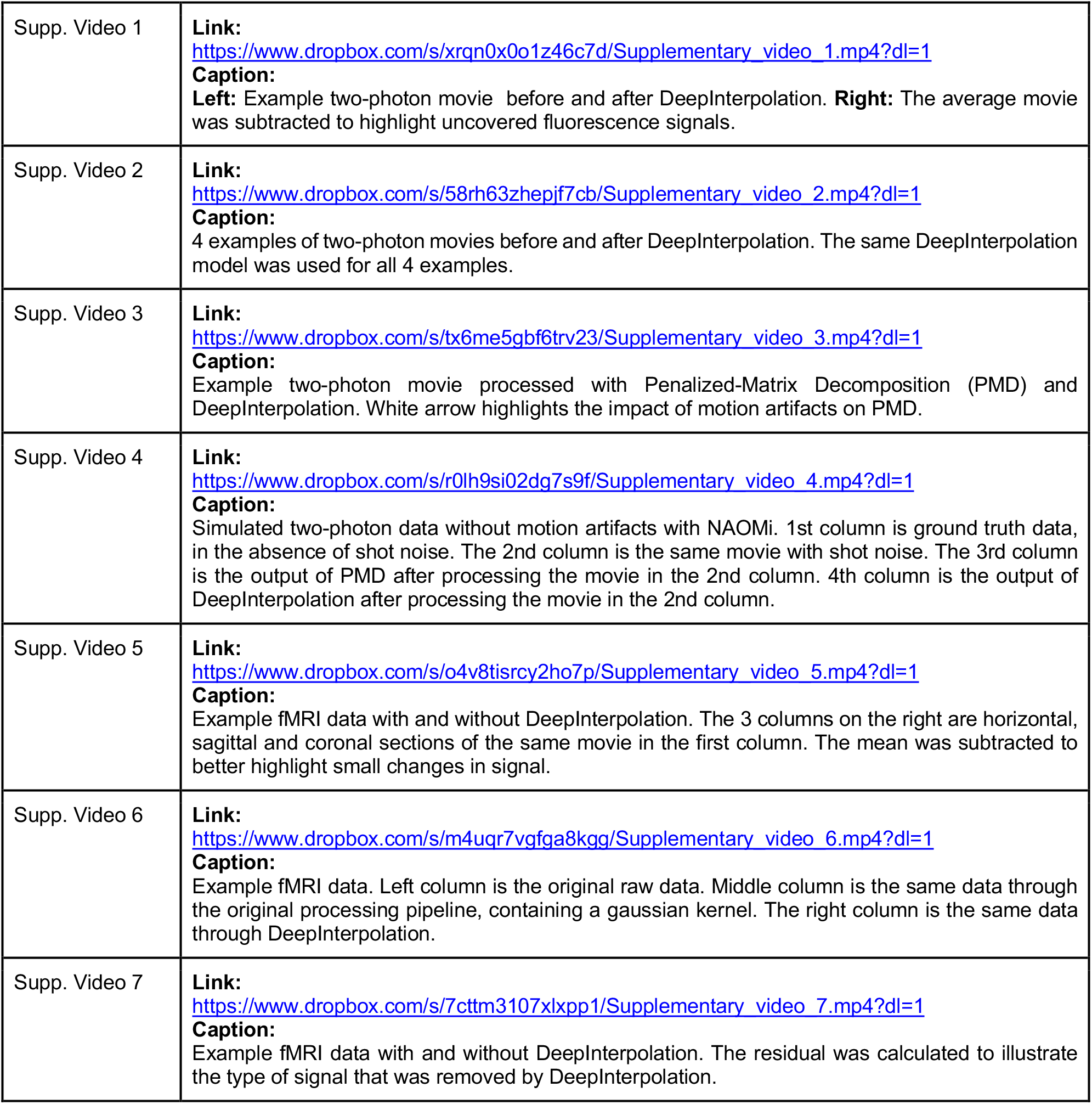

## Methods

### Description of experimental data collection of datasets used in this study

We trained 4 denoising neuronal networks in this study: one for two-photon imaging experiments using the Ai93(TITL-GCaMP6f) reporter line, one for the Ai148(TIT2L-GCaMP6f-ICL-tTA2) reporter line^26,27^, one for Neuropixels recordings using “Phase 3a” probes^28^, and one for fMRI imaging using datasets from a study published on OpenNeuro.org^29^. All raw datasets used for training are available at the time of publication through AWS S3 buckets (Amazon.com, Inc., see https://openneuro.org/datasets/ds001246/versions/1.2.1 and https://registry.opendata.aws/allen-brain-observatory/). Importantly, each denoising network is tightly coupled to the inherent properties of the datasets they were trained on (acquisition parameters, sample characteristics,…).

Calcium imaging in mice was performed with two different two-photon imaging instruments^12^ (either a Scientifica Vivoscope or a Nikon A1R MP+). Laser excitation was provided by a Ti:Sapphire laser (Chameleon Vision-Coherent) at 910 nm. 512×512 Movies were recorded at 30 Hz with resonant scanners over a 400-μm field of view.

All extracellular electrophysiology recordings were carried out with Neuropixels probes^28^ acutely inserted into the brains of awake mice. Data at each recording site was acquired at 30 kHz (spike band) and 2.5 kHz (LFP band). The spike band (which was used for denoising) had a hardware gain settings of 500× and a 500 Hz high pass filter. Each probe contained 960 recording sites, with the 374 sites closest to the tip configured for recording. Neural signals were routed to an integrated base containing filtering, amplification, multiplexing, and digitization circuitry, and were streamed via an Ethernet link to the Open Ephys GUI^30^ for online visualization and disk writing. Training data consisted of 1-hour of segments extracted from 10 different ~2.5 hour long recordings (see below for information on the specific sessions used).

All fMRI datasets were collected using 3.0-Tesla Siemens MAGNETOM Trio A Tim scanner located at the ATR Brain Activity Imaging Center. Methodology was previously described^29^. Briefly, the functional images covered the entire brain (TR, 3,000 ms; TE, 30 ms; flip angle, 80°; voxel size, 3 × 3 × 3 mm; FOV, 192 × 192 mm; number of slices, 50, slice gap, 0 mm). The dataset contained fMRI data from five subjects with 3 types of scanning sessions: “ses-perceptionTraining”, “ses-perceptionTest” and “ses-imageryTest”. We trained our denoiser on “ses-perceptionTraining” sessions and measured the denoising performance on “ses-perceptionTest” sessions.

### Visual stimulation

Visual stimuli in mice were displayed on an ASUS PA248Q LCD monitor, with 1,920 × 1,200 pixels (see original data publication for more details^12^). Stimuli were presented monocularly, and the monitor was positioned 15 cm from the eye and spanned 120° × 95° of visual space. Each monitor was gamma corrected and had a mean luminance of 50 cd/m^2^. To account for the close viewing angle, spherical warping was applied to all stimuli to ensure that the apparent size, speed, and spatial frequency were constant across the monitor as seen from the mouse’s perspective.

### Pre-processing steps

Two-photon imaging movies were motion-corrected similarly to our previous publication^12^. The motion correction algorithm relied on phase correlation and only corrected for rigid translational errors. We used these motion corrected movies for denoising.

For Neuropixels recordings, the median across channels was subtracted to remove common-mode noise. The median was calculated across channels that were sampled 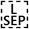simultaneously, leaving out adjacent (even/odd) channels that are likely measuring the same 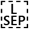spike waveforms, as well as reference channels that contain no signal. For each sample, the median value of channels N:24:384, where N = [1,2,3,…,24], was calculated, and this value was subtracted from the same set of channels. Importantly, this step removes noise that is *correlated* across channels, without affecting the independent noise targeted by DeepInterpolation. We performed denoising on the spike band after the median subtraction and offline filtering steps (150 Hz highpass) were applied.

For fMRI recordings, we used raw, unprocessed Nifti-1 volume data provided on OpenNeuro.org (see https://openneuro.org/datasets/ds001246/versions/1.2.1).

### Detection of ROIs in two-photon data

We used two different segmentation algorithms in this study. The first one was described in a previous publication and leverages a succession of morphological filters to extract binary masks surrounding active pixels^12^. These masks are publicly available through the AllenSDK (see https://allensdk.readthedocs.io/en/latest/). In all analysis related to **Supp. Fig. 5**, we applied these binary masks to both raw and DeepInterpolation movies to extract matching calcium traces.

In all analysis related to **Fig. 2**, we used the sparse mode in Suite2P^16^ to extract all individual filters. We used default parameter values of this mode except for threshold_scaling, which was set to 3. With the increased SNR achieved with DeepInterpolation, this single change limited the proportion of false positives in the final set of filters. Using Suite2p sorting GUI, we then manually sorted filters. We excluded filters present at the edge of the image, filters created by motion artifacts as well as features that did not cover any neuronal segment present in the max projection image. Somatic and non-somatic ROIs were also manually sorted based on the presence of a soma-excluded region in the filter. Filters that included a blurred out of plane soma were considered non-somatic. Once the final set of filters were extracted, we reapplied those weighted masks either to the original raw movie or the movie after DeepInterpolation to extract individual traces.

### Detection of neuronal units in electrophysiological data

Kilosort2 was used to identify spike times and assign spikes to individual units^31^. Traditional spike sorting techniques extract snippets of the original signal and perform a clustering operation after projecting these snippets into a lower-dimensional feature space. In contrast, Kilosort2 attempts to model the complete dataset as a sum of spike “templates.” The shape and locations of each template is iteratively refined until the data can be accurately reconstructed from a set of *N* templates at *M* spike times, with each individual template scaled by an amplitude, *a*.

Kilosort2 was applied to the original and denoised datasets using identical parameters (all default parameters, except for ops.Th, which was lowered from [10 4] to [7 3] to increase the probability of detecting low-amplitude units). Because the spike detection threshold is relative to the overall noise level per channel, the absolute value of the threshold was lower following DeepInterpolation.

The Kilosort2 algorithm will occasionally fit a template to the residual left behind after another template has been subtracted from the original data, resulting in double-counted spikes. This can create the appearance of an artificially high number of ISI violations for one unit or artificially high zero-time-lag synchrony between nearby units. To eliminate the possibility that this artificial synchrony will contaminate data analysis, the outputs of Kilosort2 are post-processed to remove spikes with peak times within 5 samples (0.16 ms) and peak waveforms within 5 channels (~50 microns).

Kilosort2 generates templates of a fixed length (2 ms) that matches the time course of an extracellularly detected spike waveform. However, there are no constraints on template shape, which means that the algorithm often fits templates to voltage fluctuations with characteristics that could not physically result from the current flow associated with an action potential. The units associated with these templates are considered “noise,” and are automatically filtered out based on 3 criteria: spread (single channel, or >25 channels), shape (no peak and trough, based on wavelet decomposition), or multiple spatial peaks (waveforms are non-localized along the probe axis). A final manual inspection step was used to remove any additional noise units that were not captured by the automated algorithm.

### Quality control for electrophysiological units

Units with action-potential-like waveforms detected by Kilosort2 are not necessarily high quality. To ensure that units met basic quality standards for further analysis, we filtered them using three different quality metrics, computing with the ecephys_spike_sorting Python package (github.com/AllenInstitute/ecephys_spike_sorting):

- **ISI violations score < 0.5**. This metric is based on refractory period violations that indicate a unit contains spikes from multiple neurons. The ISI violations metric represents the relative firing rate of contaminating spikes. It is calculated by counting the number of violations <1.5 ms, dividing by the amount of time for potential violations surrounding each spike, and normalizing by the overall spike rate. It is always positive (or 0), but has no upper bound. See^32^ for more details.
- **Amplitude cutoff < 0.1**. This metric provides an approximation of a unit’s false negative rate. First, a histogram of spike amplitudes is created, and the height of the histogram at the minimum amplitude is extracted. The percentage of spikes above the equivalent amplitude on the opposite side of the histogram peak is then calculated. If the minimum amplitude is equivalent to the histogram peak, the amplitude cutoff is set to 0.5 (indicating a high likelihood that >50% of spikes are missing). This metric assumes a symmetrical distribution of amplitudes and no drift, so it will not necessarily reflect the true false negative rate.
- **Presence ratio > 0.95**. The presence ratio is defined as the fraction of blocks within a session that include 1 or more spikes from a particular unit. Units with a low presence ratio are likely to have drifted out of the recording, or could not be tracked by Kilosort2 for the duration of the experiment.

Applying these quality metrics removed 54% of detected units in the original data, and 60% of units after denoising. Spike sorting after DeepInterpolation found more units regardless of the threshold used for each QC metric (**Supp. Fig. 6B**).

### Procedure for finding matched vs. new units

Following spike sorting steps, we searched for overlapping spikes between pairs of units detected before and after denoising. Spikes were considered to overlap if they had a peak occurring within ±5 channels and ms of one another. The number of overlapping spikes was used to compute three metrics, using the original spike trains as “ground truth”^33^.

- **Precision**. The fraction of denoised spikes that were also found in the original data.
- **Recall**. The fraction of original spikes that were also found in the denoised data.
- **Accuracy**. *N*_match_ / (*N*_denoised_ + *N*_original_ - *N*_match_)

Units were considered matched if they had an accuracy exceeding 0.7. Units were considered novel after denoising if they had a total precision (summed over all original units) less than 0.5. Units were considered undetected after denoising if they had a total recall (summed over all denoised units) less than 0.5.

### Training of denoising networks

We used 3 different sets of environments for training deep neural networks. Two-photon denoising networks were trained using TensorFlow 1.9.0, keras 2.2.4 and CUDA 9.0. Electrophysiology denoising networks were trained using TensorFlow 2.0 and CUDA 10.0 through their built-in Keras libraries. fMRI denoising networks were trained using TensorFlow 2.2 and CUDA 10.1.

We utilized NVIDIA Titan X, Geforce GTX 1080, and Tesla V100-SXD2 GPUs available on the Allen Institute internal computing clusters. The fMRI denoising network was trained on Amazon AWS using the p2.8×large and p3.8×large instance type depending on availability.

We used an L1 loss during training for both two-photon imaging and fMRI datasets. Electrophysiological datasets were trained with an L2 loss. All training was done with the RMSProp gradient descent algorithm implemented in keras. Two-photon denoiser was trained with a batch size of 5 so as to fit on available GPU memory. The learning rate was set to 5*10^−4^. The Neuropixels denoiser was trained with a batch size of 100, with the learning rate was set to 10^−4^. The fMRI denoiser was trained with a batch size of 10,000. The larger batch size was allowed by the smaller input-output size. The fMRI denoiser was trained with a learning scheduler, initialized at 10^−4^, dropping by half every 45 millions samples.

To facilitate training, all samples were mean-centered and normalized by a single shared value for each experiment during training. For two-photon movies, the mean and standard deviation was pre-calculated using the first 100 frames of the movie. For Neuropixels recordings, the mean and standard deviation was pre-calculated using the 200,000 samples. For fMRI recordings, we extracted the centered volume of the movie that was 1/4^th^ of the total movie size and averaged all voxels.

A detailed step by step pseudocode description of this process is available in **Supp. Appendix 1**.

### Inference of final datasets with trained networks

Once a DeepInterpolation network was learned, inference was performed by streaming entire experiments through the same fixed network. To match the conditions in training, each dataset was mean-centered and normalized following the same procedure used during training. Output denoised data were brought back to their original scale after going through the DeepInterpolation model.

### Quantification of SNR

In the two-photon data, SNR was defined as the ratio of the mean fluorescence value divided by the standard deviation along the temporal dimension. In ideal photon shot-noise limited conditions, this SNR is proportional to the square root of N, where N is the photon flux detected per pixel.

In the Neuropixels data, SNR for individual units was defined as the ratio of the maximum peak-to-peak waveform amplitude to the RMS noise on the peak channel. In the fMRI data, tSNR of individual voxels was defined as the ratio of the mean BOLD signal value divided by standard deviation along the temporal dimension.

### Analysis of natural movie responses

#### Noise and signal correlation analysis in two-photon data

For each natural movie presentation, we extracted the corresponding ROI traces. The signal correlation was computed by averaging all 10 presentations of the movie and calculating the Pearson pairwise correlation of those averaged responses between pairs of ROIs.

The noise correlation was calculated by calculating the Pearson correlation of individual trial responses between a pair of ROIs and then averaging all 10 presentations of the movie.

#### Response reliability in two-photon data

The response reliability was calculated by first calculating the pearson correlation matrix of an ROI individual response to a single presentation of a natural movie with other trials. For each ROI, we then excluded auto-correlation values and averaged out all pairwise combinations. This process yielded a single reliability measure for each ROI.

### Significant responses in Neuropixels data

When analyzing spike data, the trial-to-trial correlation of the natural movie response depends heavily on the bin size chosen. To determine responsiveness for Neuropixels data, we instead compared each unit’s natural movie PSTH (averaged across 25 trials) to a version in which all bins were temporally shuffled. A Kolmogorov-Smirnov test between the original and shuffled PSTHs was used to determine the probability that the response could have occurred by chance (*p* < 0.05 for significantly responsive units; see **Supp. Fig. 7B** for an example).

### Synthetic data generation

We created realistic synthetic calcium imaging datasets using a recent approach called in silico Neural Anatomy and Optical Microscopy (NAOMi)^14^. The code repository was kindly made available to us by Alex Song and Adam Charles. Given the computational load of the model, we generated a single dataset made of 15,000 frames simulating a 400×400 μm^2^ field of view. Except for the field of view size, all parameters were set to default values. We did not use a pre-trained network to denoise this dataset and trained our DeepInterpolation model directly on this simulated data.

### Comparison with Penalized Matrix Factorisation

A PMD docker repository was kindly made available to us by the Paninski lab and we collaborated to run the PMD algorithm in the best conditions^13^. PMD was run on an AWS instance (m5.8×large) under a pre-packaged jupyterhub environment (https://hub.docker.com/r/paninski/trefide). All movies were pre-centered and normalized using code available in the trefide package (psd_noise_estimate).

### DeepInterpolation code availability

Code for DeepInterpolation and all other steps in our algorithm are available online through a GitHub repository (https://github.com/AllenInstitute/deepinterpolation). Example training and inference tutorial code are available on the repository. Several Docker inference containers are available on Docker Hub (see list below for more details). They allow inference using the TensorFlow ModelServer framework.

Code and data to regenerate all figures presented in this manuscript is available online through a GitHub repository (https://github.com/AllenInstitute/deepinterpolation_paper).

#### Two-photon Ai93 excitatory line DeepInterpolation network

**Key recording parameters**:

- 30Hz sampling rate, 400×400 μm^2^ field of view, 512×512 pixels.
- 0.8 NA objective.
- 910 nm excitation wavelength.
- Gcamp6f calcium indicator.
- Ai93 reporter line expressed in excitatory neurons.

**Docker hub id**: 245412653747/deep_interpolation:allen_400um_512pix_30hz_ai93

#### Two-photon Ai148 excitatory line DeepInterpolation network

**Key recording parameters**:

- 30 Hz sampling rate, 400×400 μm^2^ field of view, 512×512 pixels.
- 0.8 NA objective.
- 910 nm excitation wavelength.
- Gcamp6f calcium indicator.
- Ai93 reporter line expressed in excitatory neurons.

**Pre-processing**: Individual movies were motion corrected. Each movie recording was mean-centered and normalized with a single pair of value for all pixels

**Docker hub id**: 245412653747/deep_interpolation:allen_400um_512pix_30hz_ai148

#### Neuropixels DeepInterpolation network

**Key recording parameters**:

- Neuropixels Phase 3a probes
- 374 simultaneous recording sites across 3.84 mm, 10 reference channels
- Four-column checkerboard site layout with 20 μm spacing between rows
- 30 kHz sampling rate
- 500× hardware gain setting
- 500 Hz high pass filter in hardware, 150 Hz high-pass filter applied offline

**Pre-processing**: Median subtraction was applied to individual probes to remove signals that were common across all recording sites. Each probe recording was mean-centered and normalized with a single pair of value for all nodes on the probe.

**Docker hub id**: 245412653747/deep_interpolation:allen_neuropixel

#### fMRI DeepInterpolation network

**Key recording parameters**:

- TR, 3,000 ms; TE, 30 ms; flip angle, 80°; voxel size, 3 × 3 × 3 mm; FOV, 192 × 192 mm; number of slices, 50, slice gap, 0 mm

**Pre-processing**: N/A

**Docker hub id**: 245412653747/deep_interpolation:allen_3_3_3_tr_3000_fmri

### Data availability

The two-photon imaging and Neuropixels raw data can be downloaded from the following S3 bucket: arn:aws:s3:::allen-brain-observatory

Two-photon imaging files are accessed according to Experiment ID, using the following path: visual-coding-2p/ophys_movies/ophys_experiment_<Experiment ID>.h5 We used a random subset of 1144 experiments for training the denoising network for Ai93, and 397 experiments for training the denoising network for Ai148. The list of used experiments IDs is available in json files (in deepinterpolation/examples/json_data/) on the DeepInterpolation GitHub repository. The majority of these experiment IDs are available on the S3 bucket. Some experimental data has not been released to S3 at the time of publication.

Neuropixels raw data is accessed by Experiment ID and Probe ID, using the following paths: visual-coding-neuropixels/raw-data/<Experiment ID>/<Probe ID>/spike_band.dat

The dat files have the median subtraction post-processing applied, but do not include an offline highpass filter. Prior to DeepInterpolation, we extracted 3600 seconds of data from each of the recordings listed in **Table 1**.

**Table 1.** Details of Neuropixels data used for DeepInterpolation training and inference. To train the fMRI denoiser, we used datasets that can be downloaded from an S3 bucket: arn:aws:s3:::openneuro.org/ds001246 We trained our denoiser on all “ses-perceptionTraining” sessions, across 5 subjects (3 perception training sessions, 10 runs each).

| Experiment ID | Probe ID | Start time | Brain Structures | Used for training |
| --- | --- | --- | --- | --- |
| 778998620 | 792626851 | 4665.8179 s | VISl (LM), CA1, CA3 | No |
| 787025148 | 792586842 | 4672.1967 s | VISp (V1), SUB, LP | No |
| 768515987 | 773549856 | 4672.2224 s | VISrl (RL), CA1, DG, APN | Yes |
| 794812542 | 810758787 | 4665.8332 s | VISrl (RL), CA1, DG, APN | No |
| 778998620 | 792626853 | 4665.8179 s | VISal (AL), CA1, CA3, DG, LGv | No |
| 767871931 | 773462997 | 4672.2352 s | VISal (AL), CA1, CA3, DG, LT | Yes |
| 779839471 | 792645501 | 4672.3841 s | VIS, CA1, CA3, DG, LGd | No |
| 771160300 | 773621948 | 4672.3791 s | VISal (AL), CA1, CA3, DG, LGd | Yes |
| 781842082 | 792586881 | 4672.1312 s | VISp (V1), SUB, SGN | No |
| 793224716 | 805124815 | 4675.2401 s | VISrl (RL), CA1, DG, LP, APN | No |

