## Supplementary Material for "Removing independent noise in systems neuroscience data using DeepInterpolation"

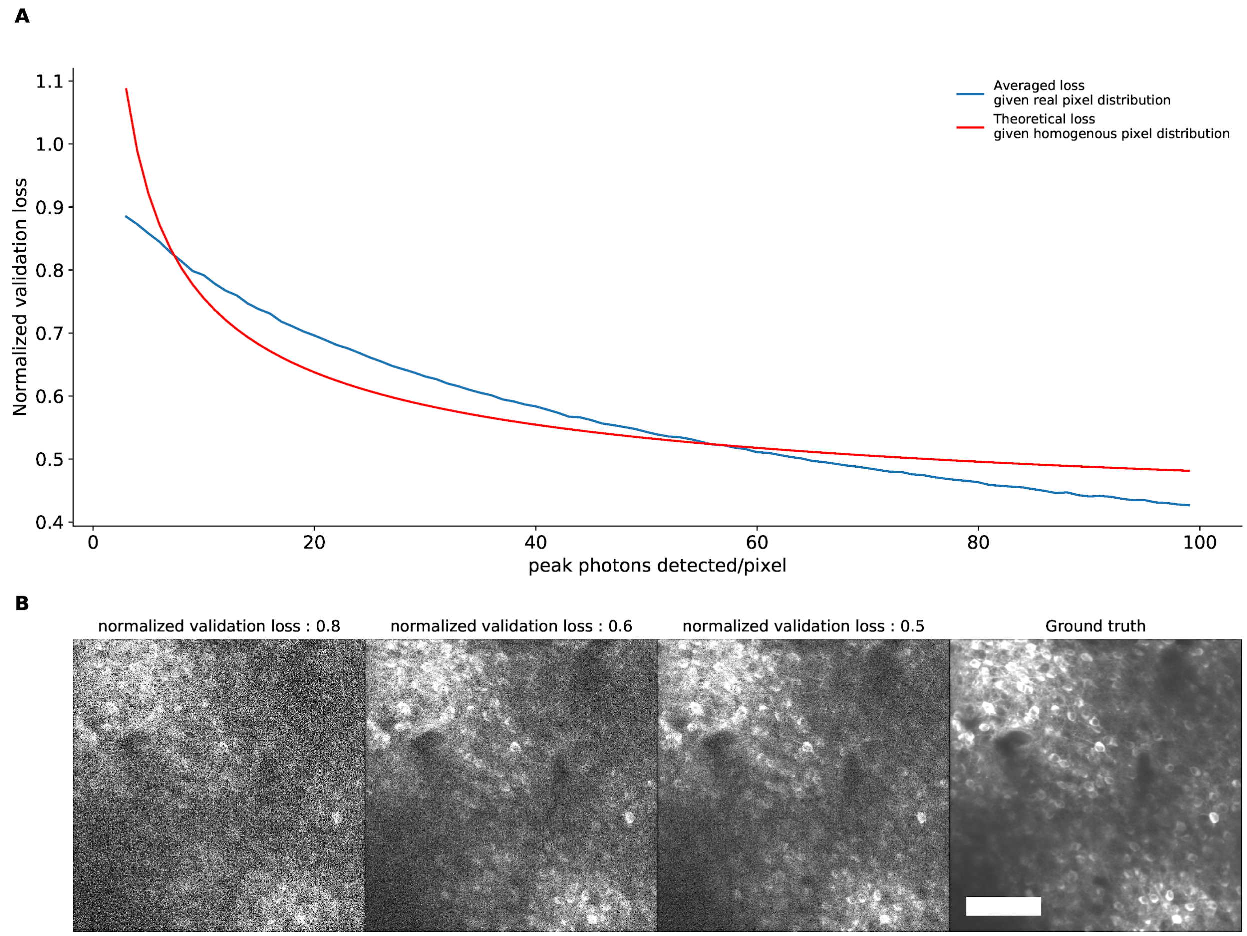


**Supp. Fig. 1 | Simulation of the relationship between mean absolute error and ground truth. (A)** The normalized mean absolute error was simulated between an averaged two photon image (Ground truth, see panel B) and various levels of shot noise. Shot noise was modulated by changing the peak number of photons detected in the image per pixel. Increasing the number of photons leads to lower errors. Denoising images contaminated with shot noise cannot yield reconstruction errors below this noise floor. As a result the reconstruction noise floor is bound by the maximum photon flux detected. Red curve was calculated assuming an homogeneous distribution of intensity and a shot noise limited SNR. (**B**) Example individual images associated with a specific normalized validation loss. See Method and Appendix I for a description of how the loss is calculated. Scale bar is 100 μm.

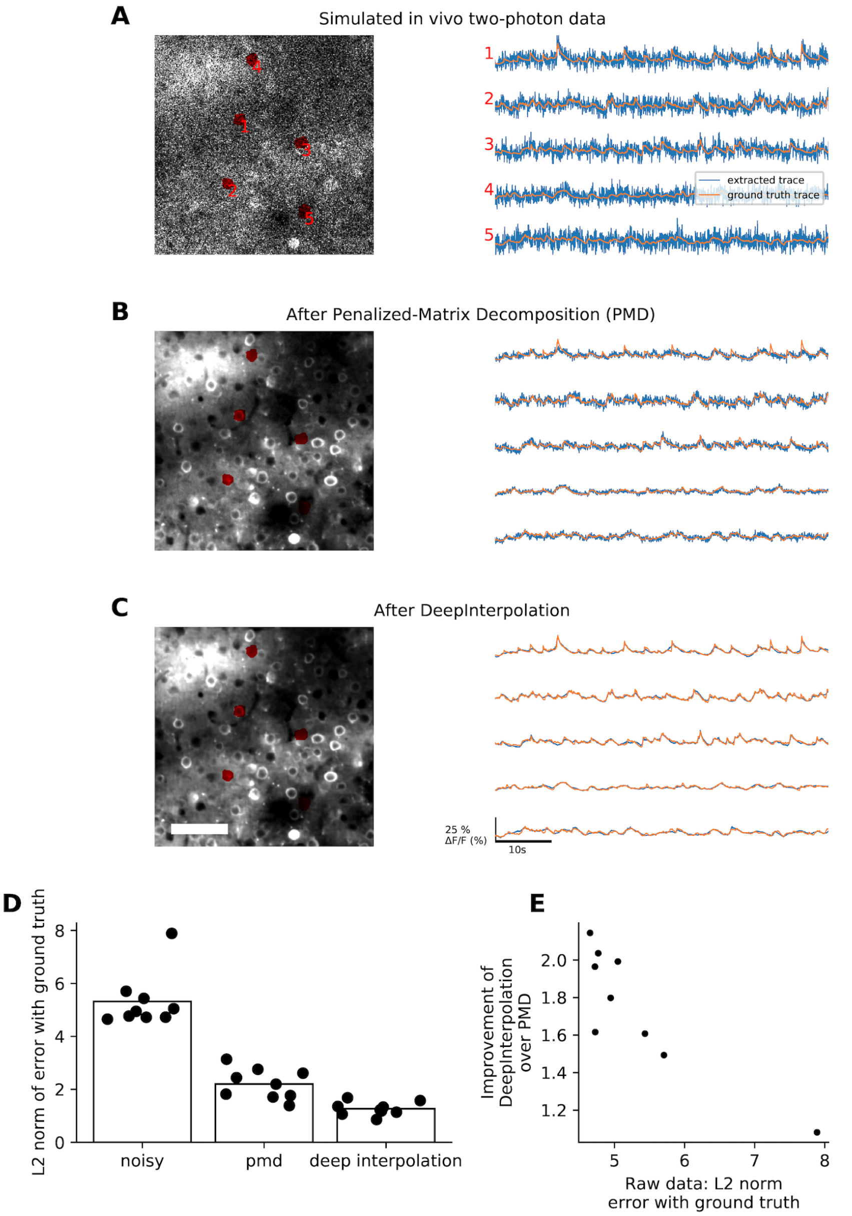


**Supp. Fig. 2 | Comparison of DeepInterpolation and Penalized Matrix Decomposition (PMD)**[1] **on simulated calcium movies with ground truth. (A)** A simulated two photon calcium movie was generated using an *in silico* Neural Anatomy and Optical Microscopy (NAOMi) simulation[2]. (**A, left**) A few ROI were manually drawn to extract the associated traces (**A, right**) of the simulated movie with (blue trace) and without sources of noise (ground truth, orange). (**B**) The same movie and traces after PMD denoising. (**C**) The same movie and traces after DeepInterpolation. Scale bar is 100 µm. (**D)** Quantification of average mean squared error (L2) between noisy traces and ground truth, as well as between PMD and DeepInterpolation with ground truth. DeepInterpolation provided the smallest reconstruction error. (**E**) Quantification of the improvement in L2 norm against the original noisy error. The cells with the smallest initial error sees the biggest gain in reconstruction.

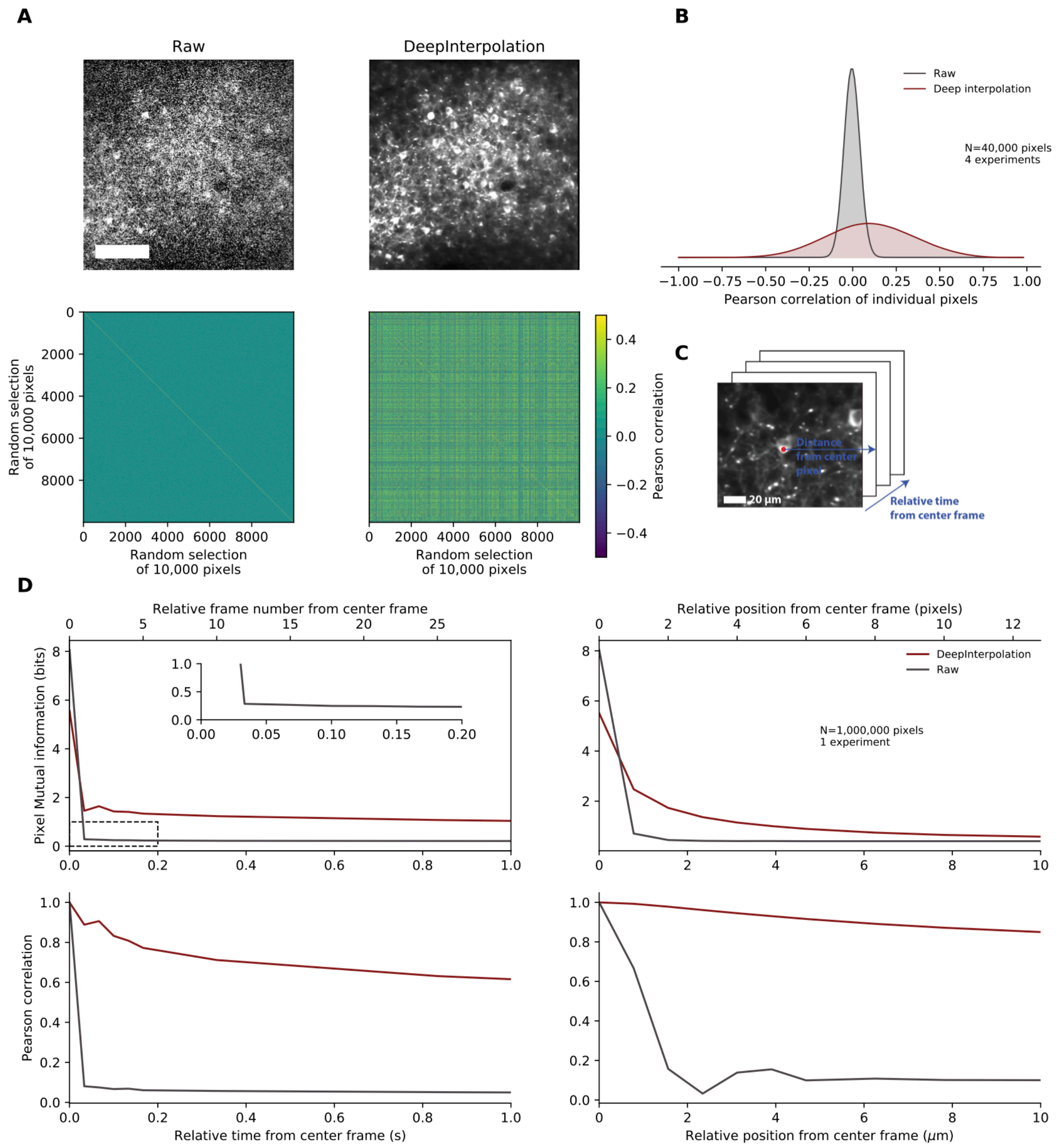


**Supp. Fig. 3 | Correlation and mutual information of pairs of pixels before and after DeepInterpolation.** (**A, top row**) Individual frames of a two photon calcium movie before and after DeepInteropolation. Scale bar is 100 µm. (**A, bottom row**) Pairwise Pearson correlation of a random selection of 10,000 pixels from top movie. **(B)** Distribution of pearson correlation for 40,000 pixels randomly selected from 4 different experiments (KS test comparing Raw with DeepInterpolation: $p$ = 9e$^{-}$

[71], $n$ = 1000 pixels randomly selected) (**C**) Schematic illustrating the spatial and temporal distances used in (D). (**D, left, top row**) Mutual information between a center pixel and consecutive frames as indicated in schematic (C). Mutual information is largely enhanced after Deep interpolation. The mutual information at the origin is the pixel entropy, illustrating how many bits of information are necessary to encode an individual pixel independently. Inset highlights a small baseline mutual information between consecutive frames in raw data. (**D, left, bottom row**). Same as (**D, top**) but plotting the pearson correlation. (**D, right, top row**). Mutual information between a center pixel and its neighboring pixel in the same frame along an horizontal axis as shown in (C ). (**D, right, bottom row**). Same as (D, top) but plotting the pearson correlation.

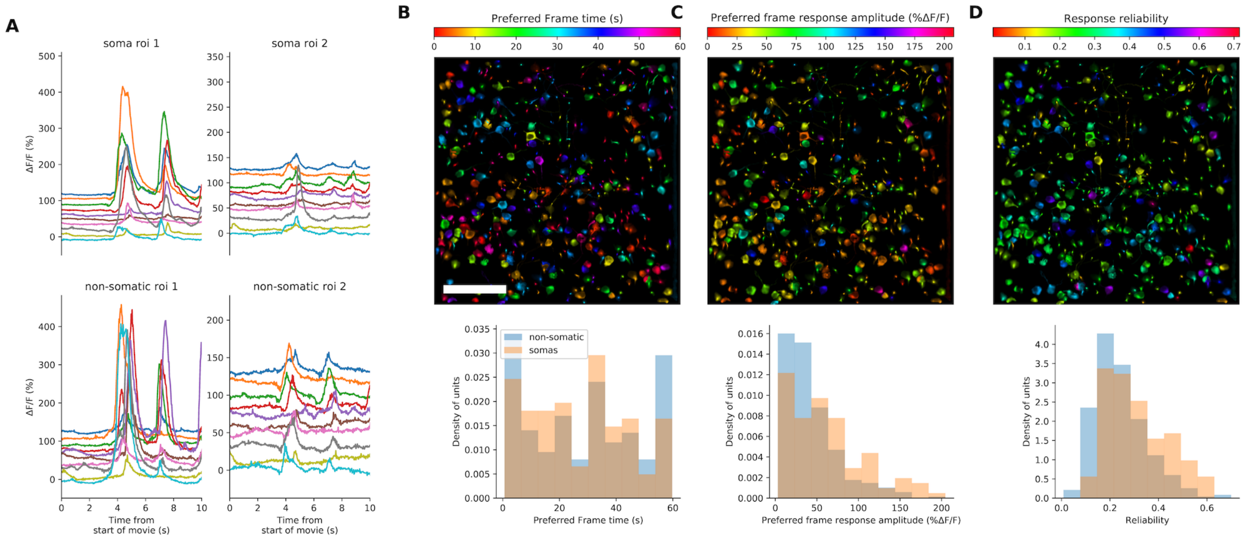


**Supp. Fig. 4 | Natural movie responses extracted from somatic and non-somatic compartments after DeepInterpolation. (A)** Example calcium responses to 10 repeats of a natural movie for 4 examples ROIs, both somatic and non-somatic **(B,C,D top)** Analysis of the sensory response to 10 repeats of a natural movie visual stimulus. Each ROI detected after DeepInterpolation was colored by the preferred movie frame time (**B**), the amplitude of the calcium response at the preferred frame (**C**) and the response reliability (**D**). Scale bar is 100 µm. **(B bottom)** Distribution preferred frame of the natural movie for both somatic and non-somatic ROIs in the same field of view. **(C bottom)** Same as (B) for response amplitude at the preferred time. (**D bottom**) Reliability of all ROIs response to the natural movie. Reliability was computed by averaging all pairwise cross-correlation between each individual trial (see Methods).

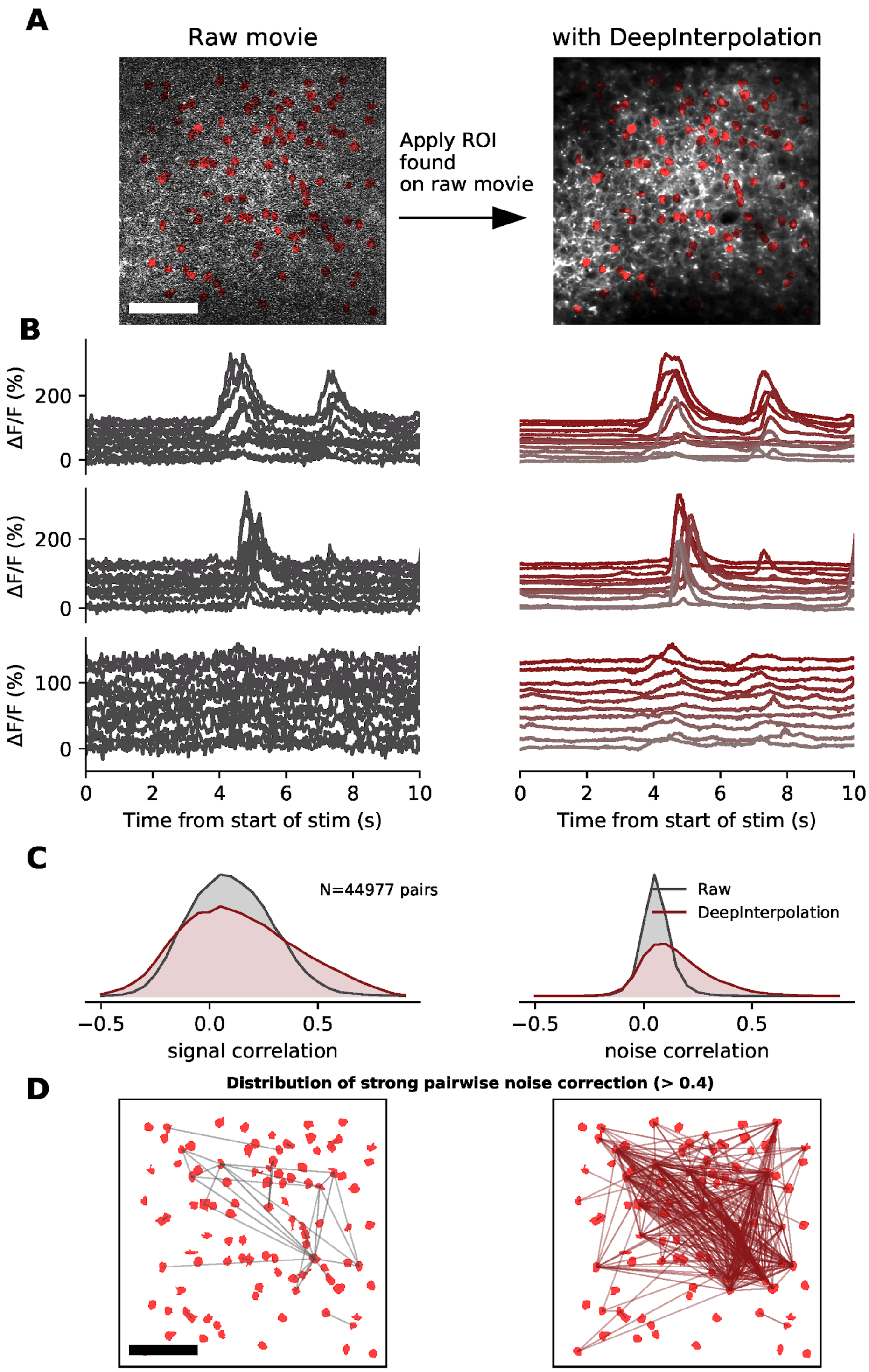


**Supp. Fig. 5 | Applying region-of-interests (ROIs) detected on noisy raw data also showcases increased signal and noise correlation. (A)** 99 ROIs detected on the original, non-denoised, movie are overlaid on top of a single frame from the original movie as well as the same frame after DeepInterpolation.

These ROI were used to extract traces in (B). Scale bar is 100 µm. (**B**) Example temporal response from 3 neuronal somas to 10 repeats (one trace per repeat) of a natural movie presentation. Left: ROI filter was applied to the original movie. Right: ROI filter was applied to the denoised movie. **(C, left)** signal correlation (average correlation coefficient between the average temporal response of a pair of neurons) for all pairs of ROI in for both raw and denoised traces. **(C, right)** noise correlation (average correlation coefficient at all time points of the mean-subtracted temporal response of a pair of neurons) for all pairs of ROI in **(A).** Distributions were created from 4 separate experiments with a total of 44,977 pairs of neurons. (**D**) Pairs of neuronal somas with high noise correlation (>0.4) are connected with a straight line for the original two-photon data (37 pairs on the left) and after DeepInterpolation (329 pairs on the right).

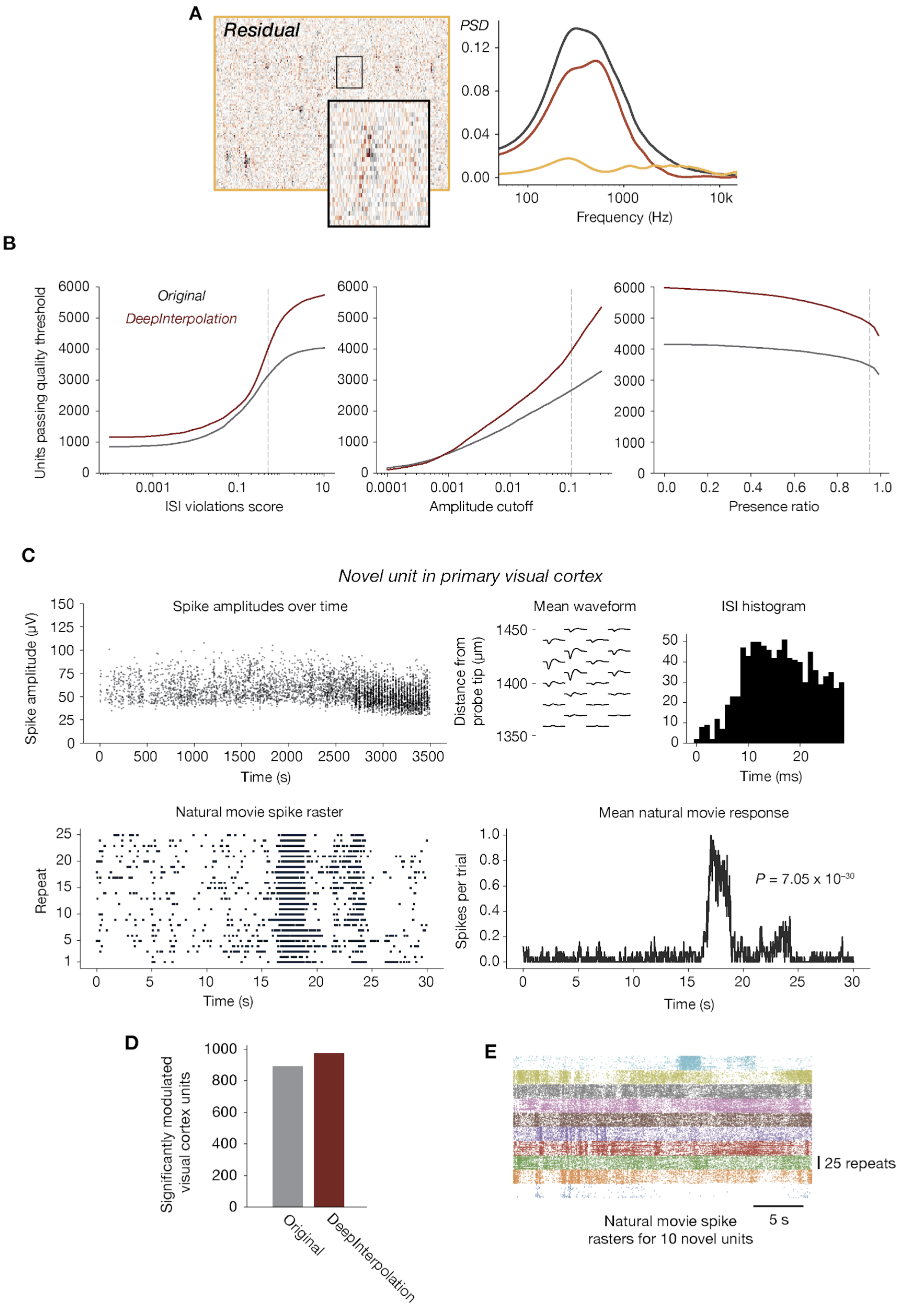


**Supp. Fig. 6 | Extended data for Neuropixels recordings.** (**A** *Left:* Heatmap of the residual (denoised subtracted from the original data), for the same data shown in Figure 3B. *Right:* Power spectra for the original (gray), denoised (red), and residual (yellow) data. **(B)** Number of units passing a range of quality control thresholds, for three different unit quality metrics. QC thresholds used in this work are indicated with dashed lines. As the ISI violations score threshold decreases (left panel), more and more contaminated units are excluded. As the amplitude cutoff threshold decreases (center panel), more and more incomplete units are excluded. As the presence ratio threshold increases (right panel), more and more unstable or artificially split units are excluded. **(C)** Physiological plots for one example unit from V1 that was only detectable after denoising (<20% of spike times were included in the original spike sorting results). This unit displays a highly reliable response to the natural movie stimulus (*P*-value calculated using the Kolmogorov–Smirnov test between the distribution of spikes per trial for the original and shuffled response). (**D**) Number of units with significant response modulation to a repeated natural movie stimulus, for all cortical units detected before and after denoising. (**E**) Raster plots for 10 high-reliability exemplar units detected only after denoising, aligned to the 30 s natural movie clip. Each color represents a different unit.

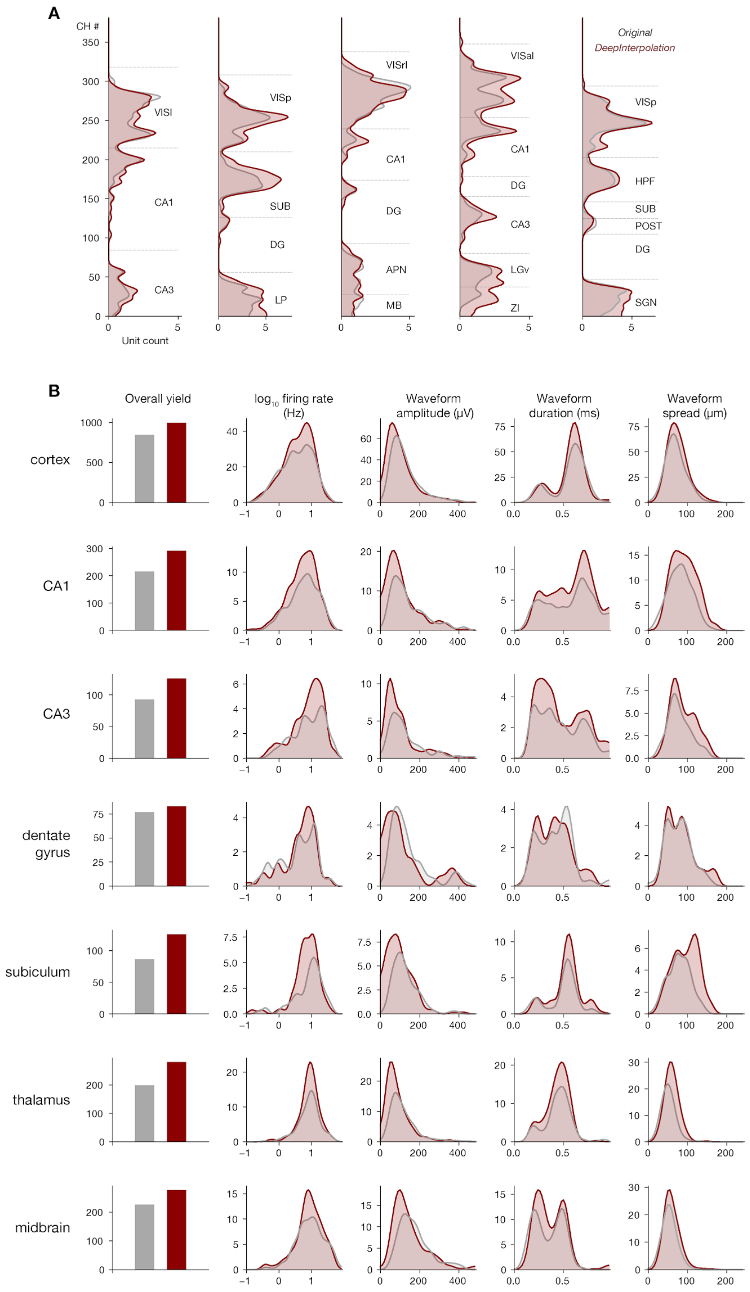


**Supp. Fig. 7 | Impact of DeepInterpolation on unit yield in different brain regions. (A)** Example unit density histograms for five example probe insertions, for original (gray) and denoised (red) data. Shaded regions represent the density of detected units passing automated QC criteria. Dashed lines represent the location of major structural boundaries. **(B)** Distributions of five unitwise metrics for seven different brain regions (see Methods for details). Histograms represent the absolute number of units before and after DeepInterpolation, aggregated across up to 10 probe insertions.

**Appendix 1 | DeepInterpolation pseudocode for denoising two-photon $Ca^{2+}$ imaging.**

1. **Training**
    a. Generate a randomized list of $N_{training}$ frames from P separate two photon movies.
    b. Calculate a single pair of sample mean and pixel standard deviation for each movie using all pixels in the first 100 frames.
    c. Store randomized list of frame index, movie data storage location and associated sample mean and standard deviation in a json file.
    d. Repeat a,b, c for validation test data on a separate set of $N_{test}$ movies.
    e. Load json file associated with training samples and test samples.
    f. Load json file with training meta-parameters.
    g. Initialize training data generators for both training and testing data. Generator is pulled from a local library of generators based on training meta-data. Training generator is directly streamed from network disk location during training by multi-threaded workers. Validation generator is pre-loaded in memory for caching.
    h. Initialize training network architecture. Architecture is pulled from a local python library based on a single string descriptor.
    i. Initialize training callbacks to save and monitor model progression throughout training.
    j. Initialize loss to "mean_absolute_error".
    k. Initialize optimizer to RMSProp.
    l. Compile model for training.
    m. Start training loop on local GPUs
        i. For each batch of training, load 5 samples from the randomized list. Both input and output samples are z-scored using pre-computed mean and standard deviation (see **b**)
        ii. Monitor training and validation loss throughout training.

iii. Save trained model every 12500 samples.

iv. Interrupt training based on validation training convergence.

**2. Inference**

a. Select one movie for inference

b. Calculate mean and standard deviation of 100 frames initial segment

c. Load trained DeepInterpolation model

d. For each frame in the movie (excluding $N_{pre}$ and $N_{post}$ frames respectively at the onset and end of the movie):

i. Z-score $N_{pre}$ and $N_{post}$ frames respectively before and after the selected frames, using pre-computed mean and standard deviation.

ii. Predict the output frame using the DeepInterpolation model

iii. Convert output frame back to original pixel values using precomputed mean and standard deviation.

iv. Save result

### References


1. Buchanan, E. K. *et al.* Penalized matrix decomposition for denoising, compression, and improved demixing of functional imaging data. doi:10.1101/334706.
2. Charles, A. S., Song, A., Gauthier, J. L., Pillow, J. W. & Tank, D. W. Neural Anatomy and Optical Microscopy (NAOMi) Simulation for evaluating calcium imaging methods. doi:10.1101/726174.
